# Integrating untargeted metabolomics, genetically informed causal inference, and pathway enrichment to define the obesity metabolome

**DOI:** 10.1101/734707

**Authors:** Yu-Han H. Hsu, Christina M. Astley, Joanne B. Cole, Sailaja Vedantam, Josep M. Mercader, Andres Metspalu, Krista Fischer, Kristen Fortney, Eric K. Morgen, Clicerio Gonzalez, Maria E. Gonzalez, Tonu Esko, Joel N. Hirschhorn

## Abstract

**Background:** Obesity and its associated diseases are major health problems characterized by extensive metabolic disturbances. Understanding the causal connections between these phenotypes and variation in metabolite levels can uncover relevant biology and inform novel intervention strategies. Recent studies have combined metabolite profiling with genetic instrumental variable (IV) analyses to infer the direction of causality between metabolites and obesity, but often omitted a large portion of untargeted profiling data consisting of unknown, unidentified metabolite signals.

**Methods:** We expanded upon previous research by identifying body mass index (BMI)-associated metabolites in multiple untargeted metabolomics datasets, and then performing bidirectional IV analysis to classify these metabolites based on their inferred causal relationships with BMI. Meta-analysis and pathway analysis of both known and unknown metabolites across datasets were enabled by our recently developed bioinformatics suite, PAIRUP-MS.

**Results:** We identified 10 known metabolites that are more likely to be the causes (e.g. alpha-hydroxybutyrate) or effects (e.g. valine) of BMI, or may have more complex bidirectional cause-effect relationships with BMI (e.g. glycine). Importantly, we also identified about 5 times more unknown than known metabolites in each of these three categories. Pathway analysis incorporating both known and unknown metabolites prioritized 40 enriched (*p* < 0.05) metabolite sets for the cause versus effect groups, providing further support that these two metabolite groups are linked to obesity via distinct biological mechanisms.

**Conclusions:** These findings demonstrate the potential utility of our approach to uncover causal connections with obesity from untargeted metabolomics datasets. Combining genetically informed causal inference with the ability to map unknown metabolites across datasets provides a path to jointly analyze many untargeted datasets with obesity or other phenotypes. This approach, applied to larger datasets with genotype and untargeted metabolite data, should generate sufficient power for robust discovery and replication of causal biological connections between metabolites and various human diseases.

## INTRODUCTION

Abnormal blood metabolite levels are important, frequent, and quantifiable feature of obesity and its associated phenotypes, which are major health problems globally^1–5^. Recently, systematic metabolite profiling (metabolomics) studies have described widespread alterations in the obesity metabolome and identified metabolite markers associated with risk of obesity-related diseases^6–9^. However, these studies broadly have two key analytic challenges limiting the biological interpretation and scope of their findings: these correlative studies have not generally been able to distinguish the cause and effect relationships between metabolites and phenotypes, and only a portion of the thousands of metabolite signals measured by untargeted profiling technology could be chemically identified and thereby routinely investigated.

Genetic instrumental variable (IV) analysis (for causal inference) and novel bioinformatics tools (for analysis of untargeted metabolite data) now provide the means to overcome these limitations and enhance our understanding of the metabolome of any phenotype. The genetic IV framework, also known as Mendelian randomization, uses genetic variants as instruments to infer causality from observational data in the presence of unmeasured confounding, provided certain methodological assumptions are met^10,11^. Bidirectional genetic IV analysis, using in turn genetic variants affecting metabolite levels and variants affecting a phenotype such as body mass index (BMI), offers a way to ascribe directionality of causal relationships and to prioritize potentially causal metabolite-phenotype associations. Previous genetic IV studies have utilized variants identified in genome-wide association studies (GWAS) to infer causality between obesity-related phenotypes and curated sets of metabolites (e.g. branched-chain and aromatic amino acids)^12–16^. However, most studies did not perform comprehensive bidirectional IV analysis and only focused on the metabolites that could be identified and curated from profiling data, thus likely capturing only a limited slice of obesity biology and, even within that constraint, not assessing causality.

Previously, metabolites of unknown chemical identities – a large portion of untargeted profiling data – were mostly excluded from analyses (including GWAS) because inter-study comparison and biological interpretation were technically onerous or intractable^17,18^. To address these issues, we recently developed a bioinformatics suite, PAIRUP-MS^18^, to match up unknown metabolites across mass spectrometry-based untargeted profiling datasets, thereby enabling meta-analysis of multiple datasets and increasing statistical power for detecting biologically interesting unknowns. In addition, PAIRUP-MS provides a framework for annotating unknown metabolites using preexisting metabolic pathways and performing pathway analysis incorporating both known and unknown metabolites.

In this study, we demonstrate how the combination of bidirectional genetic IV framework and PAIRUP-MS can be used to analyze multiple untargeted metabolomics datasets and characterize causal connections between a phenotype and the metabolome. We identified both known and unknown BMI-associated metabolites, and then performed GWAS for each metabolite and for BMI, followed by bidirectional genetic IV analysis to identify metabolites likely to be causes or effects of obesity. In addition, we highlighted distinct biological pathways enriched for the cause versus effect metabolites, confirming that the bidirectional IV approach prioritized two distinct sets of BMI-associated metabolites. This initial work illustrates an approach that can now be generalized and scaled up to much larger datasets, which will enable well-powered studies to uncover novel metabolic causes and effects of obesity or any other phenotype of interest.

## MATERIALS AND METHODS

A schematic overview of our analysis plan is shown in **Figure 1** and each step is described in more detail below.

**Figure 1.**
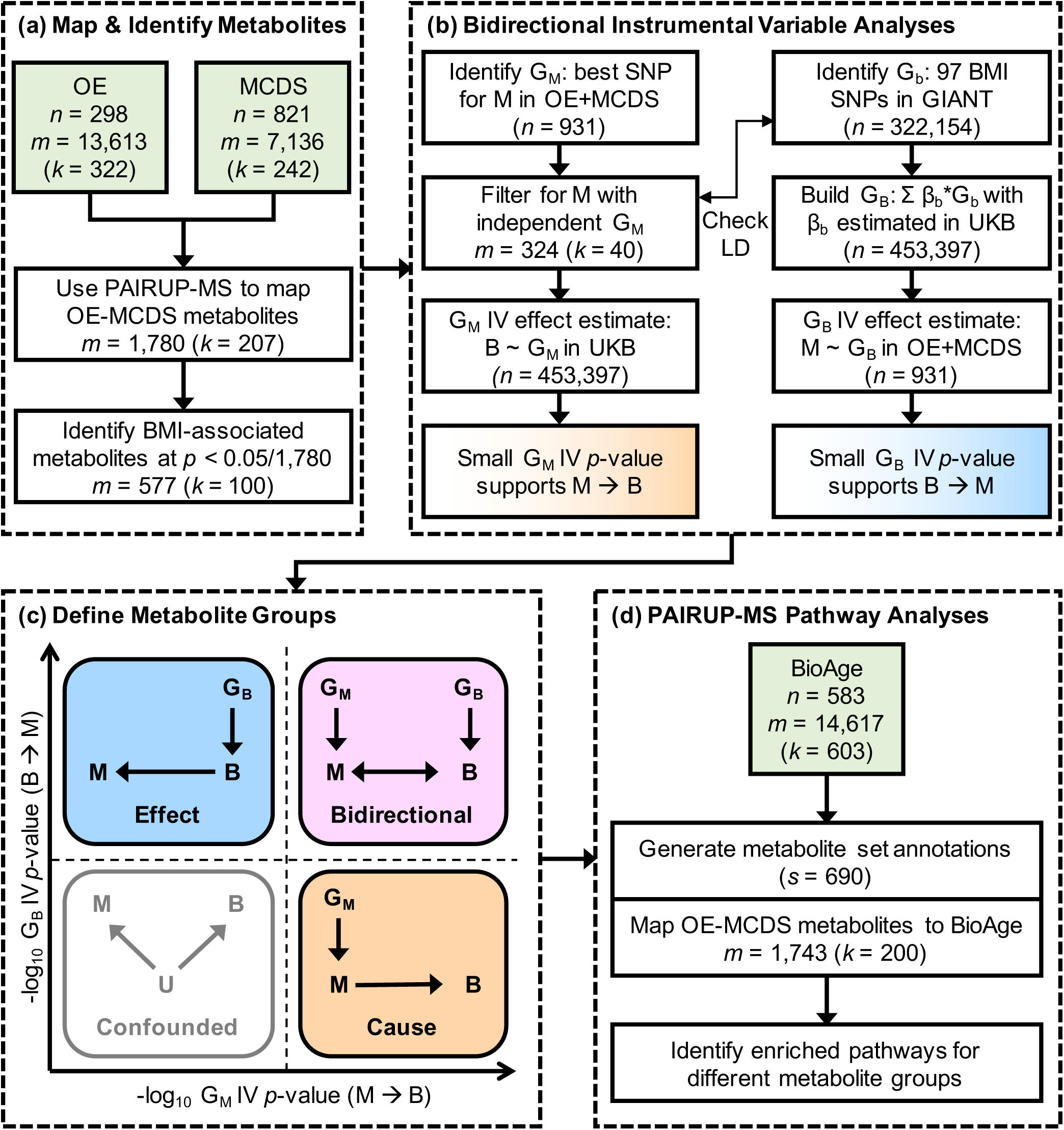
Overview for identifying and characterizing causal connections in the obesity metabolome. **(a)** OE and MCDS metabolomics datasets, matched using PAIRUP-MS, were used to identify known and unknown metabolites associated with BMI. **(b)** Independent genetic instruments (G_M_) for the BMI-associated metabolites were selected using OE and MCDS data, and then used to test for a metabolite-to-BMI (M → B) causal effect in UKB; in parallel, BMI genetic instrument (G_B_), a polygenic risk score built using GIANT BMI-associated SNPs (G_b_) and UKB effect estimate weights (β_b_), was used to test for a BMI-to-metabolite (B → M) causal effect in OE and MCDS. **(c)** A subset of metabolites was categorized into “cause”, “effect”, and “bidirectional” groups based on the significance of the G_M_ and G_B_ IV effect estimate *p*-values, reflecting different types of causal connections between the metabolites and BMI. **(d)** Pathway analyses of the three metabolite groups were performed using metabolite set annotations generated using PAIRUP-MS and BioAge data. Y ∼ X, regression of Y on X; U, unmeasured confounder; *n*, number of samples; *m*, number of metabolites; *k*, number of known (or shared known) metabolites; *s*, number of metabolite sets.

### Metabolomics datasets and data processing

#### Study populations

The study populations have been described previously^18–20^: (1) Obesity Extremes (OE): N = 300 sampled equally from lean, obese, and the general Estonian Biobank (EB) population, (2) Mexico City Diabetes Study (MCDS): N = 865 in a prospective study, and (3) BioAge Labs Mortality Study (BioAge): N = 583 in a retrospective mortality study nested in EB. All participants provided informed consent. Individual studies were approved by their respective local ethics committees. Boston Children’s Hospital Institutional Review Board approved this research.

#### Metabolite data processing

Untargeted liquid chromatography-mass spectrometry (LC-MS) profiling of plasma samples, quality control, and missing value imputation of the data have been described previously^18^. The processed OE dataset contained 298 samples and 13,613 metabolite signals (322 known); MCDS contained 821 samples and 7,136 signals (242 known); BioAge contained 583 samples and 14,617 signals (603 known). Within each dataset, we performed rank-based inverse normal transformation on each signal and used the resulting abundance *z*-scores in downstream analyses. For OE and MCDS data used in BMI and genetic association analyses, we performed covariate adjustment (age, sex, and fasting time for OE; age and sex for MCDS) before the transformation. In this paper, we refer to both known and unknown metabolite signals as “metabolites” for simplicity, recognizing that an unknown signal does not always represent an independent, functional circulating metabolite.

### Mapping and identifying BMI-associated metabolites (Figure 1a)

#### Mapping metabolites across datasets

Using the imputation-based matching algorithm in PAIRUP-MS^18^, we identified 1,780 metabolite pairs (207 shared known metabolites measured in both datasets and 1,573 matched unknown or unshared known metabolites) that could be compared directly across OE and MCDS and restricted subsequent analyses to these metabolites. For pathway analyses requiring the BioAge-based metabolite set annotations (see below), we furthered mapped 1,743 (200 shared known and 1,543 matched) of these metabolite pairs to metabolites measured in BioAge.

#### Identifying BMI-associated metabolites

Within each cohort, we adjusted raw BMI (available for 298 OE and 818 MCDS samples) for age and sex, performed rank-based inverse normal transformation on the residuals, and used the resulting BMI *z*-scores in all further analyses. (Since the OE obese and lean samples were drawn from the BMI extremes of EB, all EB samples were used to calculate population-based *z*-scores.) To identify BMI-associated metabolites, we performed linear regression of BMI on each metabolite within each dataset, followed by inverse variance weighted meta-analysis across the two datasets, and applied a Bonferroni significance threshold (*p* < 0.05/1,780) in the meta-analysis.

### Bidirectional instrumental variable analyses (Figure 1b)

#### Metabolite instrument (G_M_) selection

GWAS of the BMI-associated metabolites using 294 OE and 637 MCDS samples (with available genetic data) and subsequent inverse variance weighted meta-analysis were performed as described previously^18^. To select G_M_, we first identified the SNP (single nucleotide polymorphism) with the best meta-analyzed *p*-value for each metabolite. Next, to avoid using redundant G_M_, we “clumped” the best SNPs for all metabolites to select independent SNPs that have *r*^2^ < 0.5 or are > 250 kb apart, and only kept the independent SNPs as G_M_ (along with their best-associated metabolites) in further analyses. For known metabolites in our causality groups (see below), we performed an additional sensitivity analysis using (where available) genome-wide significant (*p* < 5 × 10^−8^) SNPs from published metabolite GWAS^21–26^ as individual G_M_.

#### BMI instrument (G_B_) selection

We used 97 BMI-associated SNPs (G_b_) previously identified in GIANT^27^ and their effect estimates (β_b_) in our UK Biobank (UKB) GWAS to calculate a weighted genetic risk score for use as G_B_ (i.e. G_B_ = Σ β_b_ × G_b_). We performed BMI GWAS in UKB using 453,397 European-ancestry samples and sex-combined BMI *z*-scores, using BOLT-LMM^28^ to account for relatedness and population structure (**Supplementary Text 1**). Analysis of the UKB data was approved by its governing Research Ethics Committee and the Broad Institute Institutional Review Board. The GIANT, UKB, and metabolomics cohorts have no known sample overlap. We confirmed that G_B_ was significantly associated with BMI in OE and MCDS and that none of the G_b_ are in linkage disequilibrium (*r*^2^ > 0.3) with the selected G_M_.

#### Testing for metabolite-to-BMI causal effect using G_M_

The association between BMI and each G_M_ was extracted from the UKB GWAS summary statistics and used to calculate the Wald ratio IV effect estimate of the metabolite (shared known or matched pair) on BMI. The *p*-value for the Wald estimate was calculated using an asymptotic standard error estimate as described previously^29^. This *p*-value – a test of the null hypothesis of no causal effect of the metabolite – was used to rank metabolites as more or less likely to be causal for BMI.

#### Testing for BMI-to-metabolite causal effect using G_B_

We performed linear regression of each BMI-associated metabolite on G_B_ in OE and MCDS separately, followed by inverse variance weighted meta-analysis. The Wald ratio IV effect estimate of BMI on each metabolite was calculated using the meta-analyzed statistics, and the corresponding *p*-value was used to rank metabolites as more or less likely to be effects of BMI. As a sensitivity analysis, we performed the MR-PRESSO global test^30^ to assess overall horizontal pleiotropy among the individual SNPs (G_b_) contained within G_B_, using metabolite-G_b_ association in the OE-MCDS meta-analysis and BMI-G_b_ association in UKB for 96 of 97 BMI SNPs (rs2033529 was excluded due to absence in our metabolite GWAS).

### Defining cause, effect, and bidirectional metabolite groups (Figure 1c)

To rank BMI-associated metabolites as more or less likely to be the causes or effects of obesity, we used the −log_10_ *p*-value of the IV effect estimate for either the metabolite (G_M_) or BMI (G_B_) instrument, reasoning that the statistical significance of these estimates is informative. Metabolites in the top and bottom quartiles of these two *p*-value-based rankings were assigned to three distinct groups corresponding to different types of causal connections with BMI: (1) “cause”: metabolites that were ranked in the top quartile using G_M_ and the bottom using G_B_, and thus are likely to be upstream causes for BMI; (2) “effect”: metabolites that were ranked in the bottom quartile using G_M_ and the top using G_B_, and thus are likely to be downstream effects of BMI; (3) “bidirectional”: metabolites that were in the top quartiles of both rankings, suggesting complex bidirectional cause-effect relationships with BMI.

### Pathway analyses of the defined metabolite groups (Figure 1d)

#### Metabolite set annotations

BioAge data and PAIRUP-MS were used to generate metabolite set annotations as described previously^18^. Briefly, pathway annotations from ConsensusPathDB^31^ were consolidated into 690 metabolite sets with unique metabolite combinations (i.e. one metabolite set may correspond to multiple pathways containing identical sets of metabolites). We then used metabolite correlations in BioAge to expand the metabolite sets to include both known and unknown metabolites, calculating a membership score for each metabolite in each set.

#### Pathway analyses

We applied the pathway analysis framework in PAIRUP-MS to identify enriched metabolite sets for the cause, effect, and bidirectional metabolite groups we defined. We compared each of the three groups individually versus all other BMI-associated metabolites and, in a fourth analysis, compared the cause versus effect groups. First, for each metabolite set in each comparison analysis, a two-tailed Wilcoxon rank-sum test was performed to compare the membership scores of the two groups of metabolites. Next, to account for correlation structure in our data, iterations of this procedure were performed using “null” metabolite groups to calculate a permutation-based enrichment *p*-value for each metabolite set (**Supplementary Figure 1**).

### Performing *m*/*z* query for unknown metabolites

To assess if the unknown metabolites captured information redundant to the known metabolites in our dataset (and to look up potential identities of unknowns classified in the three causality groups), we performed *m*/*z* query as described previosly^18^, using the “LC-MS Search” tool in the Human Metabolome Database (HMDB)^32^. The unknowns were annotated as an *m*/*z*-matched adduct of a known metabolite in our data, an *m*/*z*-matched adduct of an HMDB metabolite not identified in our data, or as a metabolite without a match in HMDB.

## RESULTS

### Identifying known and unknown metabolites associated with BMI

We used untargeted metabolomics data from OE and MCDS to identify metabolites associated with BMI. First, we identified 207 pairs of shared known metabolites measured in both cohorts, and used PAIRUP-MS to match 1,573 additional pairs of unknown or unshared known metabolites likely to represent identical or highly correlated metabolites. Then, by performing meta-analysis of both the shared known and matched pairs across the cohorts, we identified 577 BMI-associated metabolites at Bonferroni significance (*p* < 0.05/1,780), the majority of which were unknown metabolites: 418 (72.4%) consisted of two paired unknown metabolites, 59 (10.2%) consisted of a known metabolite matched to an unknown metabolite, and only 100 (17.3%) consisted of shared known metabolites. When we clustered these metabolites, we observed metabolite clusters that consisted mostly or entirely of matched pairs of unknown chemical identities (**Supplementary Figure 2**). Therefore, including these unknown metabolites in downstream analyses increased the number of candidate metabolites by nearly five-fold, and allowed us to investigate aspects of obesity biology not represented by the curated, known metabolites.

### Identifying metabolites more likely to be causal for BMI

Before we could determine whether the BMI-associated metabolites are likely to be causal for BMI, we first needed to identify the SNP best-associated with each metabolite to use as genetic instrument (G_M_ in **Figure 1**). We therefore performed GWAS of metabolite levels in both OE and MCDS, followed by meta-analysis. We identified genome-wide significant (*p* < 5 × 10^−8^) SNPs for 204 (35 shared known and 169 matched) of the BMI-associated metabolites (**Figure 2**); 66 (14 shared known and 52 matched) of these were also significant after correction for multiple hypothesis testing (*p* < 5 × 10^−8^/577). Overall, the matched, unknown metabolites showed comparable degree of genetic associations as the shared known metabolites, even in loci not associated with any of the knowns. Analyzing the unknowns thus greatly improved our ability to obtain significant and novel genetic instruments for metabolite signals, despite a relatively small GWAS sample size.

**Figure 2.**
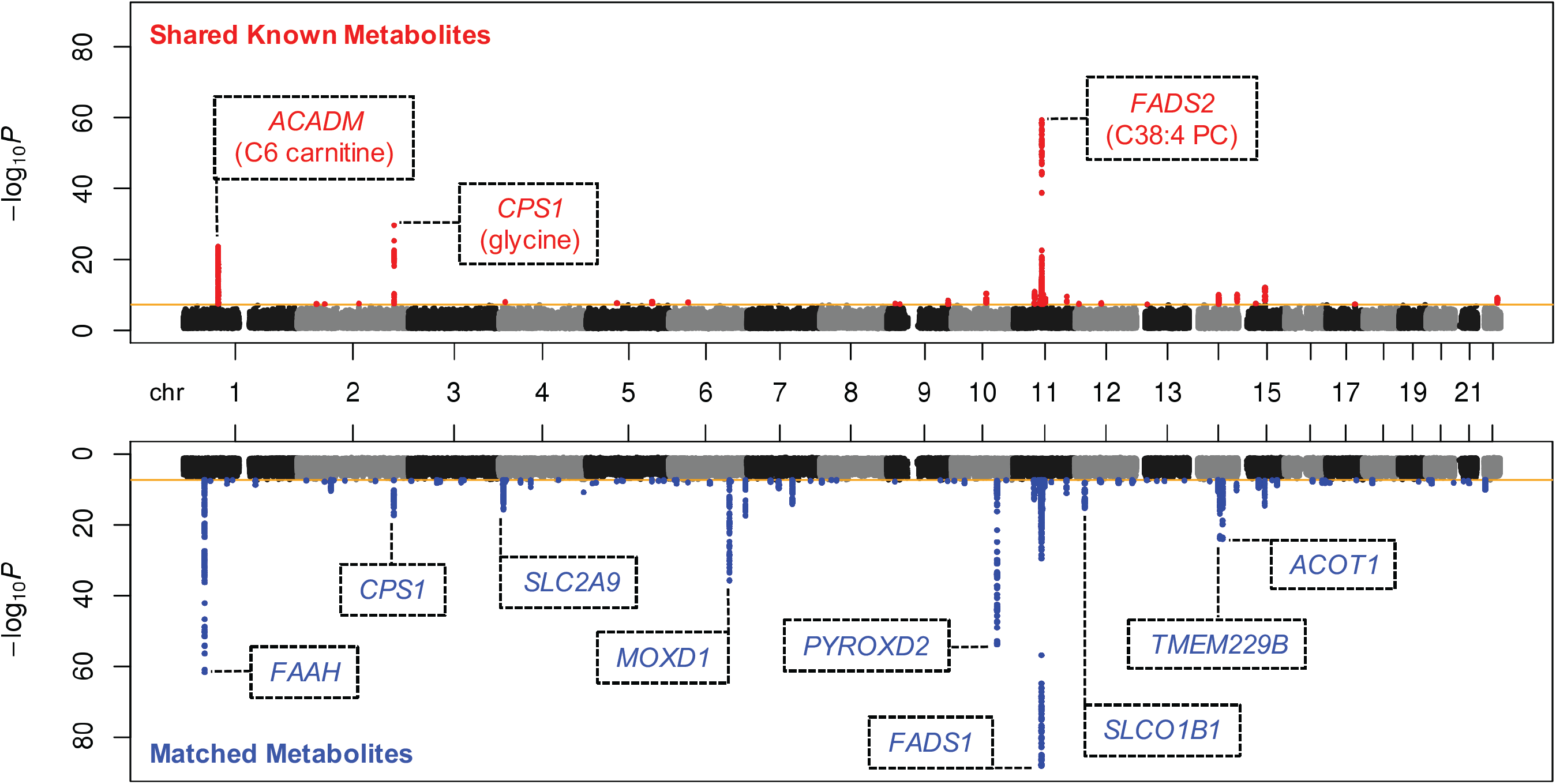
Joint Manhattan plots summarizing GWAS of BMI-associated metabolites in OE and MCDS. Genetic associations for 100 shared known (top) or 477 matched (bottom) BMI-associated metabolites were consolidated to plot the best *p*-value for each SNP (i.e. only the *p*-value for the best associated metabolite was plotted for each SNP). Genome-wide significance threshold (*p* < 5 × 10^−8^) is marked by the orange lines. Genome-wide significant SNPs are plotted in red or blue, for shared known or matched metabolites, respectively. Lead SNPs of the most significant loci *(p <* 1 × 10^−15^) are annotated with nearest genes (within 5kb), along with the best associated known metabolites if applicable.

We observed that all 577 BMI-associated metabolites had best-associated SNPs with at least suggestive significance (maximum *p* = 2.5 × 10^−6^) and therefore considered the best-associated SNP for each metabolite as potential instrument. To avoid analyzing metabolites sharing the same instruments, we included only genetically independent G_M_ (*r*^2^ < 0.5 or > 250kb apart) and the 324 (40 shared known and 284 matched) metabolites best-associated with these instruments in subsequent IV analyses (**Supplementary Table 1**). For each metabolite, we estimated the association between G_M_ and BMI using a large independent cohort, UKB, in a two-sample design to calculate the metabolite-to-BMI IV effect estimate. We identified 50 (11 shared known and 39 matched) metabolites with nominally significant (*p* < 0.05) metabolite-to-BMI IV *p*-values, which indicates that they are more likely to be upstream causes for BMI (**Supplementary Table 1**).

### Identifying metabolites more likely to be effects of BMI

Next, to determine if the BMI-associated metabolites are likely to be effects of BMI, we combined 97 BMI SNPs previously identified in GIANT into a weighted genetic risk score using UKB effect estimates as weights. As expected, the score is a valid genetic instrument for BMI (G_B_ in **Figure 1**) in OE and MCDS (meta-analyzed BMI-G_B_ association *p* = 5.9 × 10^−7^). For each of the 324 BMI-associated metabolites, we estimated the association between G_B_ and the metabolite using OE and MCDS data to calculate the BMI-to-metabolite IV effect estimate. A total of 56 (8 shared known and 48 matched) metabolites had nominally significant (*p* < 0.05) BMI-to-metabolite IV *p*-values and thus are more likely to be downstream effects of BMI (**Supplementary Table 1**).

### Defining cause, effect, and bidirectional metabolite groups

In order to further characterize the causal relationships between BMI and its associated metabolites, we ranked the metabolites based on the significance of their G_M_ and G_B_ IV effect estimate *p*-values (i.e. metabolite-to-BMI or BMI-to-metabolite IV *p*-values, respectively), and classified a subset of them into “cause”, “effect”, or “bidirectional” groups using quartile cutoffs of the rankings (**Figure 3**). We defined 25 metabolites as more likely to be cause (5 shared known and 20 matched), 26 as more likely to be effect (3 shared known and 23 matched), and 19 as more likely to be bidirectional (2 shared known and 17 matched) with respect to BMI. The shared known metabolites in each group are listed in **Table 1**; the top cause, effect, and bidirectional metabolites are alpha-hydroxybutyrate, valine, and glycine, respectively. Details for all metabolites in each group are shown in **Supplementary Table 1**. We also performed *m/z* query in HMDB to obtain potential identities for the unknowns in the matched metabolite pairs (**Supplementary Table 2**) and found only 6 out of the 60 matched pairs to be potentially redundant with the known metabolites curated in our data. Hence, we identified about 5 times more matched, unknown metabolites in the three causality categories compared to only analyzing the known metabolites. In addition, we performed sensitivity analyses to assess how our genetic IV and classification scheme would be influenced by weak instrument or pleiotropy bias (**Supplementary Text 2** and **Supplementary Table 3**); we obtained results that generally support the robustness of our approach.

**Table 1.** BMI-associated known metabolites classified into cause, effect, or bidirectional groups using the bidirectional IV effect estimate *p*-values. Y ∼ X, regression of Y on X; B, BMI; M, metabolite; covariate adjustment for B and M as described in Methods; SNP, hg19 chromsome:position is shown; β, effect size estimate; EA, effect allele (i.e. metabolite level-increasing allele). Nominally significant *p*-values (< 0.05) for IV effect estimates are in bold italic.

| Group | Metabolite | Observational Association | | Metabolite Instrument | | | $G_M$ IV Estimate ($M \rightarrow B$) | | $G_B$ IV Estimate ($B \rightarrow M$) | |
| --- | --- | --- | --- | --- | --- | --- | --- | --- | --- | --- |
| | | $\beta$ $B \sim M$ | $P$ $B \sim M$ | SNP | EA | $P$ $M \sim G_M$ | $\beta$ | $P$ | $\beta$ | $P$ |
| Cause | alpha-hydroxybutyrate | 0.235 | 7.09E-13 | 11:119745598 | T | 4.82E-07 | -0.040 | ***1.24E-03*** | -0.036 | 8.36E-01 |
|  | C34:4 PC | 0.153 | 3.45E-06 | 3:182171263 | A | 1.42E-07 | -0.027 | ***9.90E-03*** | 0.019 | 9.15E-01 |
|  | C6 carnitine | 0.207 | 2.66E-10 | 1:76224010 | C | 2.88E-24 | -0.010 | ***3.09E-02*** | -0.076 | 6.69E-01 |
|  | C18:1 CE | -0.215 | 5.54E-11 | 20:38984849 | T | 3.44E-07 | -0.016 | ***4.23E-02*** | -0.018 | 9.17E-01 |
|  | cotinine | -0.169 | 3.10E-07 | 10:123918365 | C | 1.07E-06 | 0.012 | 8.03E-02 | -0.022 | 9.02E-01 |
| Effect | valine | 0.445 | 1.20E-47 | 14:32724292 | A | 7.01E-08 | -0.003 | 6.94E-01 | 0.708 | ***1.46E-03*** |
|  | C22:6 LPC | -0.180 | 4.38E-08 | 18:71068347 | A | 3.26E-07 | -0.001 | 9.52E-01 | -0.450 | ***2.25E-02*** |
|  | C18 carnitine | -0.193 | 4.65E-09 | 6:110760008 | A | 1.27E-07 | 0.001 | 9.41E-01 | -0.351 | ***6.53E-02*** |
| Bidirectional | glycine | -0.308 | 7.55E-22 | 2:211540507 | A | 2.35E-30 | 0.030 | ***6.03E-10*** | -0.712 | ***1.59E-03*** |
|  | tyrosine | 0.376 | 6.22E-33 | 6:111477887 | C | 1.12E-07 | -0.022 | ***8.77E-03*** | 0.334 | 7.46E-02 |

**Figure 3.**
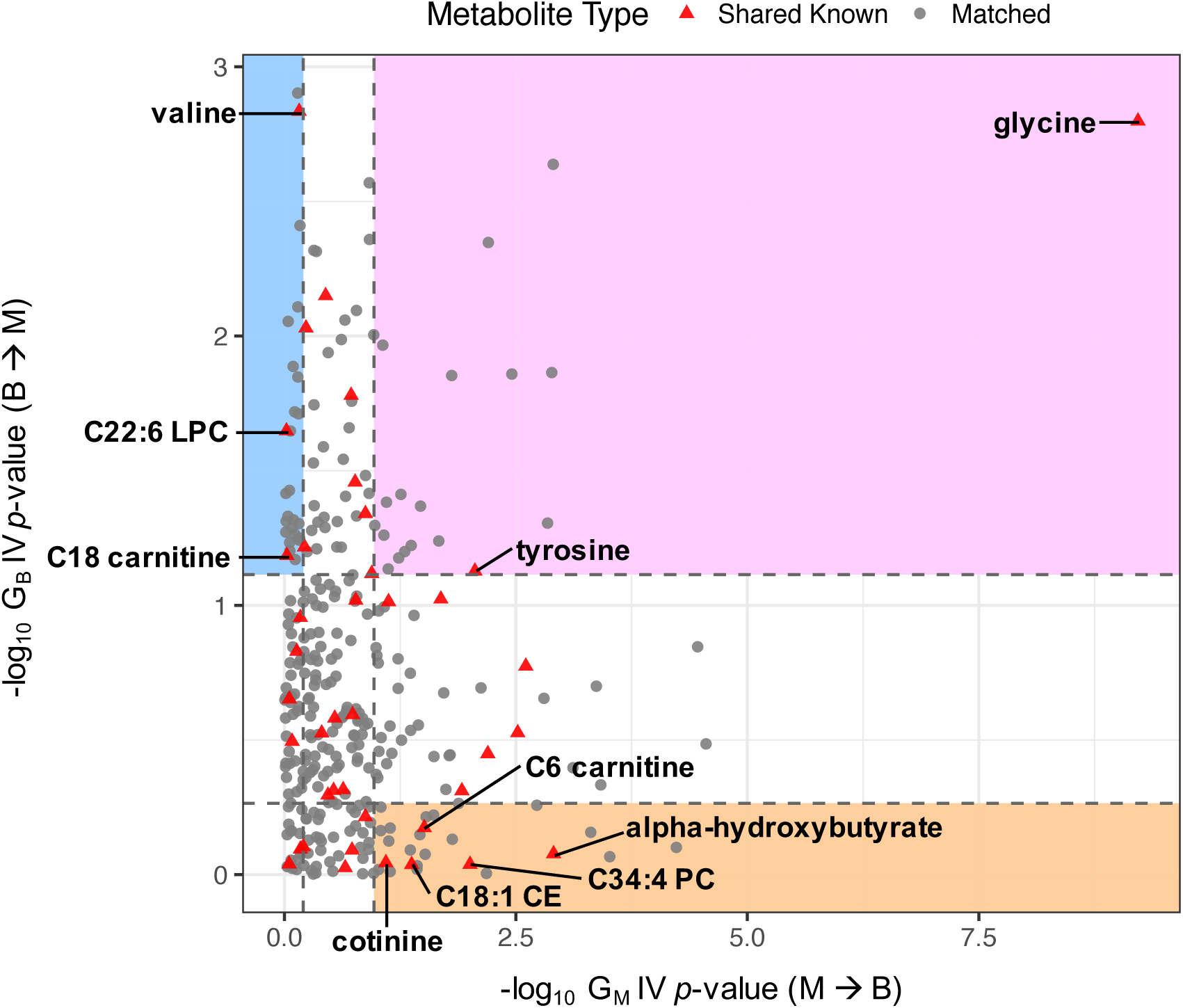
Classifying BMI-associated metabolites using IV effect estimate *p*-values for G_M_ (metabolite-to-BMI direction, x-axis) and G_B_ (BMI-to-metabolite direction, y-axis). Top and bottom quartile cutoffs along each axis are shown as dashed lines. Shared known metabolites in “cause” (orange), “effect” (blue), and “bidirectional” (pink) regions are labeled with their names.

### Prioritizing enriched pathways for cause, effect, and bidirectional metabolites

We identified many more matched, unknown metabolite pairs in the cause, effect, and bidirectional groups compared to the shared known metabolites, but it is difficult to hypothesize on their roles in obesity biology without knowing their chemical identities. Therefore, to extract useful information from the unknowns and to gain clues about the biology broadly captured by the three causality groups, we performed PAIRUP-MS pathway analyses encompassing both known and unknown metabolites, using metabolite set annotations generated from a separate cohort, BioAge. First, we carried out three separate analyses to identify pathways with nominally significant (*p* < 0.05) enrichment for metabolites in the cause, effect, or bidirectional groups, respectively, when compared against all other BMI-associated metabolites (**Supplementary Table 4**). While the most enriched metabolite sets in each analysis are associated with different pathways, several metabolite sets were enriched in multiple analyses (e.g. “NAD *de novo* biosynthesis” was enriched for both cause and effect metabolites).

Hence, in order to identify pathways that are the most distinct between the defined metabolite groups, we next performed a pathway analysis directly comparing the cause versus effect metabolites, prioritizing 40 metabolite sets at nominal significance (*p* < 0.05; **Supplementary Table 4**). The 13 cause metabolite sets (in which cause metabolites have higher membership scores than effect metabolites) are associated with various pathways, such as those connected to inflammation (e.g. nitric oxide signaling), redox metabolism (e.g. cysteine/methionine metabolism), and appetite regulation (e.g. endocannabinoid signaling). The 27 effect metabolite sets also contain varied pathways including those related to lysine catabolism, neurobiology (e.g. addiction and catecholamine biosynthesis), and stress response (e.g. FoxO signaling). While the known metabolites in our analysis have been linked to some of the enriched metabolite sets in literature, the unknown metabolites contributed most of the data used to prioritize these sets.

Finally, to better visualize the distinguishing features between the cause versus effect metabolites in terms of their roles in biological pathways, we constructed a heat map of the metabolites’ membership scores in the enriched metabolite sets using unsupervised clustering (**Figure 4**). The metabolites formed two major clusters consisting of metabolites that are mostly in the cause or effect groups, with a handful of metabolites clustering with the contrasting group (i.e. cause metabolite “misclassified” in the effect cluster or vice versa). Even more strikingly, the cause and effect metabolite sets formed two pure clusters consisting of all cause or all effect sets. This clustering pattern provides further evidence that the cause and effect metabolites we defined are involved in distinct biological processes and thus may be associated with BMI through different mechanisms.

**Figure 4.**
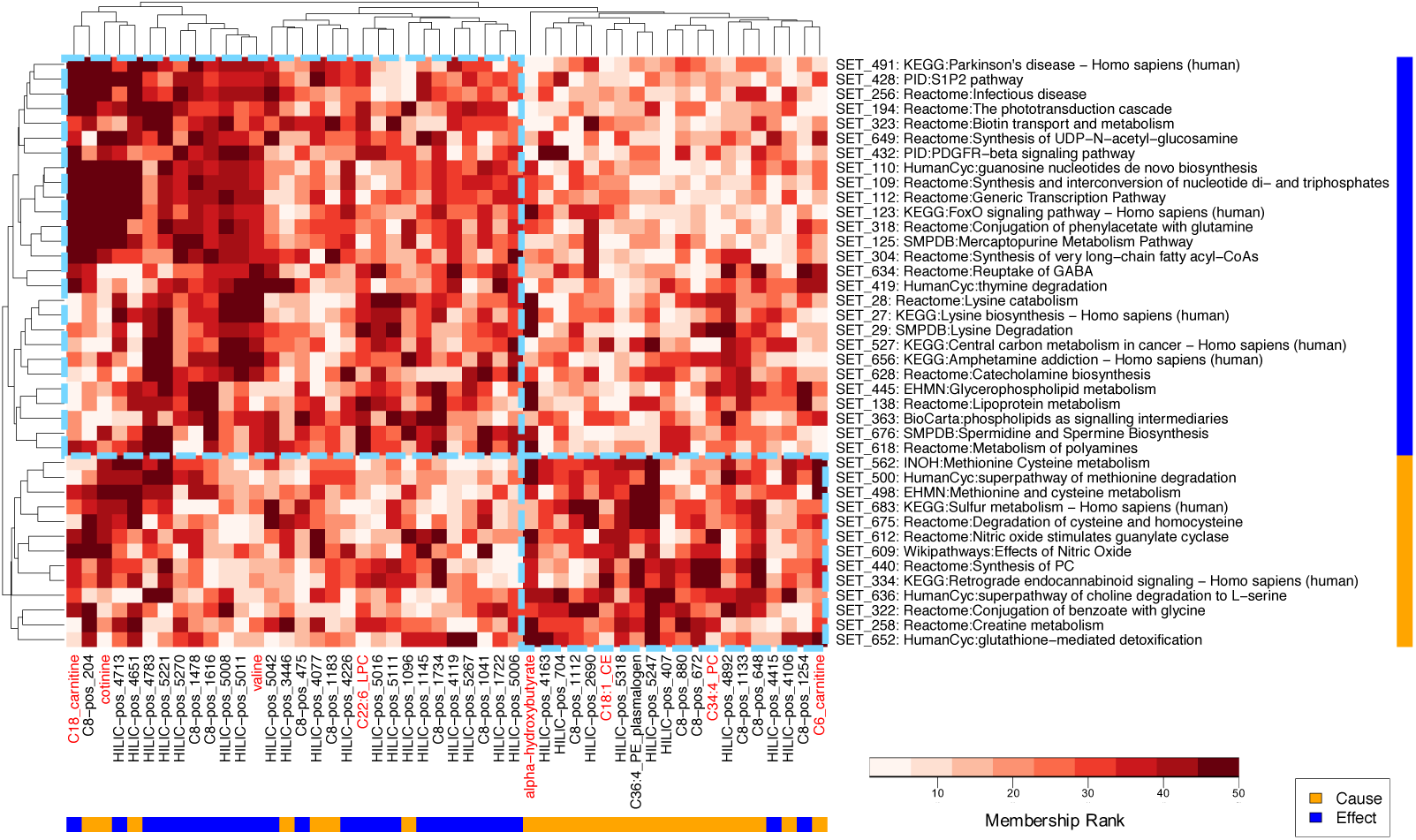
Clustered heat map of cause and effect metabolites’ memberships in metabolite sets prioritized by pathway analysis. Hierarchical clustering was performed using membership scores in the BioAge-based metabolite set annotations. Each column is a shared known (red label) or matched (black label) metabolite from the cause (yellow bar) or effect (blue bar) metabolite group. Each row is a significant (*p* < 0.05) metabolite set in pathway analysis in either the cause (yellow bar) or effect (blue bar) direction (with representative pathway name shown in label; see **Supplementary Table 4** for full pathway list). Larger number in membership rank (darker red) indicates higher membership score. Dashed light blue boxes highlight the two major cause and effect clusters according to the clustering dendrograms.

## DISCUSSION

The study of comprehensive metabolite profiles defines an exciting frontier in human pathophysiology. However, metabolite-phenotype associations discovered in metabolomics studies are often correlative in nature and additional causal inference approaches, such as genetic IV analysis, are required to help assess causality between metabolites and phenotypes. Furthermore, unknown metabolite signals are often filtered out prior to analysis of untargeted metabolomics data, greatly limiting investigation to *a priori* candidate metabolites, reducing the search space, and hindering downstream analyses such as pathway enrichment. Here we present a paradigm for combining untargeted metabolomics, genomics, and our recently described bioinformatics suite, PAIRUP-MS, to overcome these challenges. Using obesity as an exemplar state of metabolic dysregulation, we illustrate the potential utility of this approach to advance our understanding of causal connections in metabolic diseases.

In this study, we meta-analyzed hundreds of unknown metabolites from two cohorts using PAIRUP-MS, identifying novel associations between the unknowns, BMI, genetic variants, and biological pathways. Indeed, using bidirectional genetic IV analysis, we discovered about 5 times as many unknown than known metabolites with potential causal connections to BMI. While these unknowns are likely not all fully independent and functional circulating molecules, their associations with genetic variants and BMI, distinct from those with known metabolites, suggest that a sizable number of unknown metabolites reflect aspects of BMI biology not captured by known metabolites. Furthermore, the much larger number of candidate metabolites allowed us to perform PAIRUP-MS pathway analyses that account for potential redundancy, prioritizing biological pathways specific to the metabolites with cause or effect relationships to BMI. Because of the relatively small sample sizes of our cohorts, some of our results did not meet stringent multiple hypothesis testing significance thresholds; nevertheless, they demonstrate a useful and generalizable analytic framework to probe the metabolome of obesity and other diseases as larger datasets become available.

We identified novel metabolites that may be causes of obesity, as well as replicating two known metabolites, valine and tyrosine, that may be the effects of BMI^14^. The strongest causal evidence among known metabolites was for alpha-hydroxybutyrate, which has been linked to insulin resistance, oxidative stress, glutathione biosynthesis, and mitochondrial dysfunction^6,33,34^. The oxidative stress and glutathione connections are especially intriguing since “glutathione-mediated detoxification” emerged as a significant causal pathway when we compared the cause and effect metabolite groups in pathway analysis. It is also notable that the IV effect estimate of alpha-hydroxybutyrate on BMI is protective while the observational association suggests this metabolite is obesogenic. We postulate that a mitochondrial dysfunction/altered redox state linked to high alpha-hydroxybutyrate level could lead to decreased weight gain, while shared common causes, such as an obesogenic diet, may lead to increases in both alpha-hydroxybutyrate level and BMI. This example highlights the advantage of genetic IV analyses over observational studies alone to explore the potential impact of a theoretical intervention targeted to obesity-associated metabolites that have yet to be fully characterized^35,36^.

The validity of genetic IV analysis rests upon several key assumptions. Specifically, the genetic instrument must explain variation in the exposure variable and the instrument must not be associated with the outcome variable except through its relationship with the exposure (no genetic pleiotropy). Weak instrument bias towards the null and pleiotropy bias away from the null may lead to misclassification of metabolites in our three causality groups. To address weak instrument bias for our known metabolite instruments, we performed sensitivity analysis using stronger instruments from published metabolite GWAS, showing that our results are generally robust against weak instrument bias, although some misclassification is possible due to limited power of our internal instruments. However, we could not conduct similar analysis for the unknown metabolite instruments since there is not yet a straightforward way to obtain external instruments for comparison. To address pleiotropy bias for our BMI instrument, we used a recently developed method, MR-PRESSO, to show that our BMI IV estimates are likely robust against extreme cases of pleiotropy bias. We could not examine pleiotropy in the metabolite instruments due to the lack of multiple instruments for each metabolite (especially for the unknowns where additional instruments could not be obtained from published GWAS).

Larger GWAS of both known and unknown metabolites, conducted across multiple datasets, will make it possible to extend our paradigm to understand causal biological mechanisms for various metabolic diseases and alleviate the limitations described above. With more candidate metabolites and genetic instruments emerging from better-powered studies, our approach can be expanded to mediation analyses^37^, to pathway Mendelian randomization^38^, or to metabolite IV subsetting according to predicted biological pathway memberships^39^. In conclusion, this study showcases the benefit of combining untargeted metabolomics with a bidirectional genetic IV approach to define the metabolome of a major human disease state, obesity. We therefore advocate for broader sharing of untargeted metabolomics and genetic datasets, similar to the approach taken by international efforts to optimize GWAS of many other phenotypes. Broader sharing would improve power and reliability of methodological frameworks such as the one presented here, and would enable a fuller realization of the potential of metabolomics to generate important insights into human diseases.

## ACKNOWLEDGEMENTS

We thank the Broad Metabolomics Platform and SIGMA T2D Consortium for sharing data resources. This research has been conducted using the UK Biobank Resource under Application Number 11898. This work was supported by the National Heart, Lung, and Blood Institute grant F31HL126581 (Y.H.H.), National Institute of Diabetes and Digestive and Kidney Diseases grants T32DK110919 (Y.H.H.), K12DK094721 (C.M.A), and R01DK075787 (J.N.H.), Endocrine Scholars Award (C.M.A.), Doris Duke Charitable Foundation grant 215205 (J.N.H.), Estonian Research Council grants IUT20-60 (A.M.), PUT1665 (K.Fischer), and PUT1660 (T.E.), European Union through Horizon 2020 grant 692145 (A.M.), and European Union through the European Regional Development Fund Project No. 2014-2020.4.01.15-0012 (A.M.).

