## Supplementary Figure 1 for "Integrating untargeted metabolomics, genetically informed causal inference, and pathway enrichment to define the obesity metabolome"

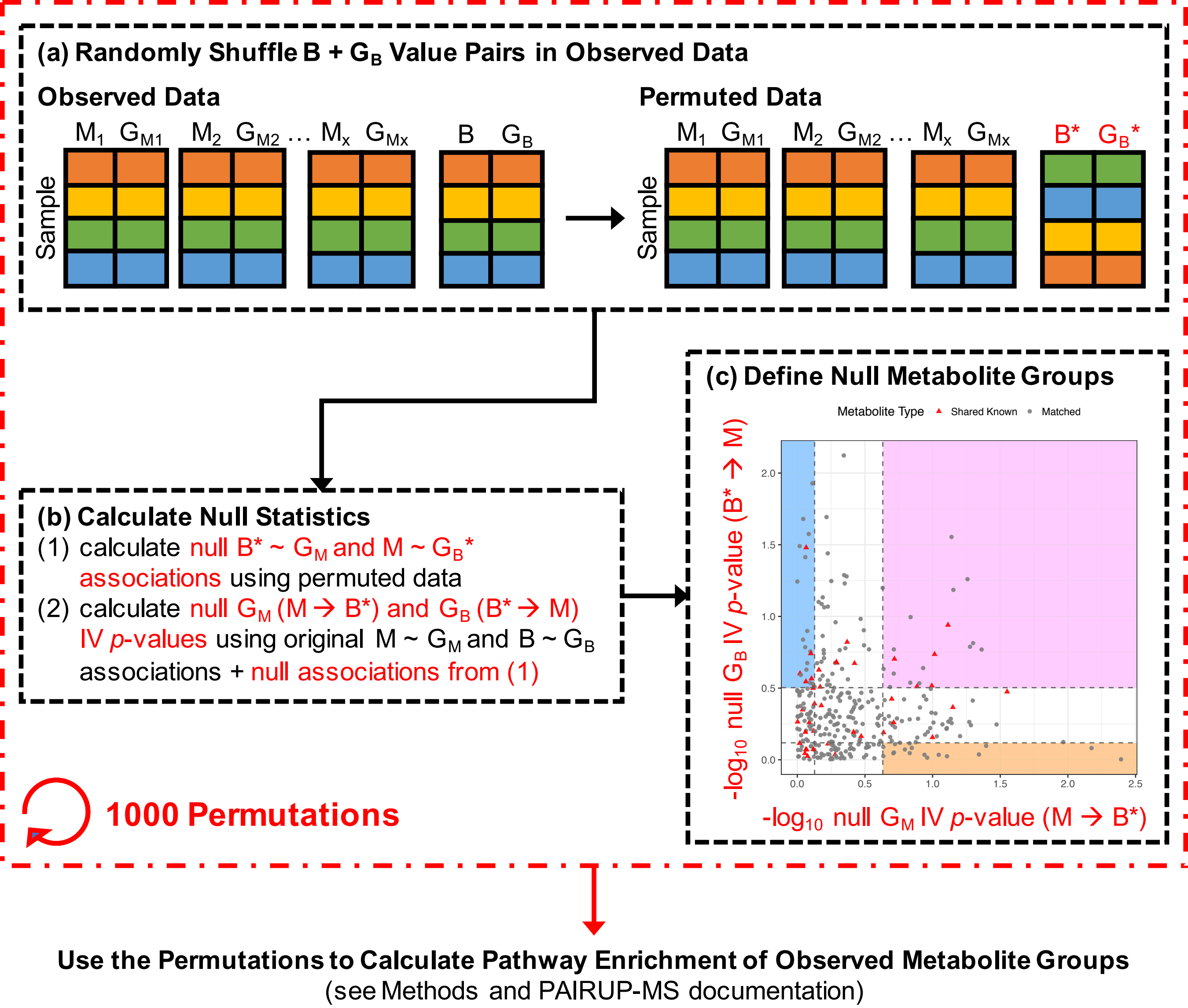


**Supplementary Figure 1. Scheme for generating null metabolite groups used in pathway analyses. (a)** For each permutation, we randomly shuffled the BMI (B) and $G_B$ value pairs in OE and MCDS data, while leaving the metabolites (M) and $G_M$ data unchanged. Same color shading in the cartoon illustration indicates data that originated from the same sample in observed data. B* and $G_B$ *, permuted BMI and $G_B$ data, respectively. **(b)** After permuting the data, we calculated null association statistics for B* ~ $G_M$ and M ~ $G_B$*, and then combined these results with original, non-permuted M ~ $G_M$ and B ~ $G_B$ statistics to calculate null metabolite-to-BMI (M → B*) and BMI-to-metabolite (B* → M) Wald ratio IV effect estimate *p*-values. Y ~ X, regression of Y on X. **(c)** We used the top and bottom quartile cutoffs of the null IV *p*-values to define null cause, effect, and bidirectional metabolite groups (mirroring how we defined the observed metabolite groups). The entire permutation procedure (steps (a) – (c)) was repeated 1000 times to generate 1000 sets of null metabolite groups, which were used to calculate the permutation-based pathway enrichment *p*-values in PAIRUP-MS pathway analyses (i.e. permutation permutation *p*-value = proportion of null rank-sum *p*-values smaller than or equal to the observed rank-sum *p*-value).
