## Supplementary Figure 2 for "Integrating untargeted metabolomics, genetically informed causal inference, and pathway enrichment to define the obesity metabolome"

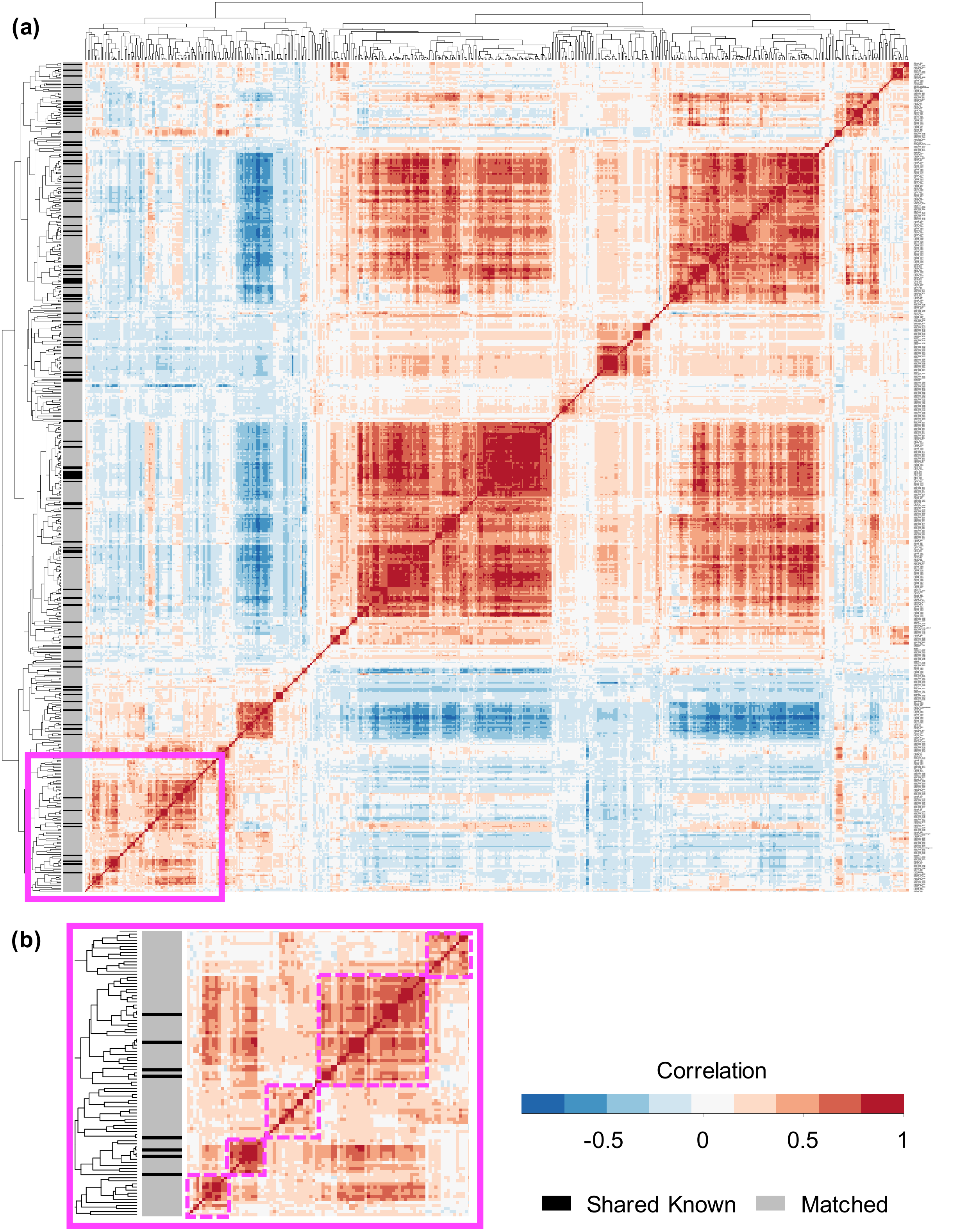


**Supplementary Figure 2. Clustered correlation heat maps of the 577 BMI-associated metabolites in OE and MCDS. (a)** Heat map of all BMI-associated metabolites. Hierarchical clustering was performed using pairwise correlations of the metabolites in OE+MCDS samples. Shared known (black) and matched (grey) metabolites are annotated by side bar on the left. **(b)** Zoomed-in view of a portion of the heat map (solid pink box in (a)), with distinct clusters (as defined by the clustering dendrograms) highlighted by pink dashed boxes.
