## Supplementary Text 1 and 2 for "Integrating untargeted metabolomics, genetically informed causal inference, and pathway enrichment to define the obesity metabolome"

**Supplementary Text 1. Details of BMI GWAS in UKB**

GWAS was performed using 453,397 European samples as determined by both genetic and self-reported ancestry. Quality control of the samples have been described previously^1^. Individuals flagged by UKB as having excess heterozygosity, excess missingness, an inferred versus genetic sex mismatch, putative aneuploidy, or currently pregnant were additionally excluded. Average BMI from up to three assessment center visits of the male and female samples were separately adjusted for covariates (average age in months, average age in months squared, genotyping platform, assessment year, and the first ten genetic principal components), followed by inverse normal transformation to calculate BMI *z­*-scores. BOLT-LMM^2^ was used to perform linear mixed model GWAS on sex-combined BMI *z*-scores.

**Supplementary Text 2. Sensitivity analyses of the metabolite classification scheme**

To assess how sensitive our genetic IV analysis and classification scheme would be to weak instrument and pleiotropy bias, we evaluated external metabolite instruments and performed a test for pleiotropy for the BMI instrument. First, to address weak instrument bias that could be caused by our G_M_, which were selected using relatively small GWAS, we searched previously published GWAS to obtain external instruments for the 10 known metabolites classified in our causality groups. Eight out of the 10 known metabolites have genome-wide significant (*p* < 5 × 10^-8^) published instruments (**Supplementary Table 3**). Four of these metabolites have published instruments in the same loci (i.e. < 100 kb) as our G_M_, indicating that a good portion of our G_M_ is replicable in larger cohorts. When looking at only the top published instruments (SNP with smallest *p*-value) for the cause and bidirectional metabolites, in all 5 cases, the metabolites showed consistent direction of causal effect compared to results obtained using our G_M_ (**Supplementary Table 3A**). Similarly, when we looked at top published instruments for the effect metabolites (i.e. metabolites that lack causal metabolite-to-BMI evidence in our data), 2 out of 3 metabolites had no significant causal effect on BMI (metabolite-to-BMI IV *p* > 0.05), also agreeing with results from our G_M_ (**Supplementary Table 3B**). The exception is valine, whose top instrument yielded a very significant causal effect estimate (metabolite-to-BMI IV *p* = 2.8 × 10^‑7^) and would reclassify valine as a bidirectional metabolite. However, there are also other valine instruments that agree with our G_M_, indicating there is heterogeneity. Overall, IV analysis results derived using previously published metabolite instruments generally support the robustness of the results obtained using our internal G_M_, despite these G_M_ being selected from a relatively small GWAS dataset.

Next, to estimate horizontal pleiotropy among individual SNPs (G_b_) contained in G_B_, we performed the MR-PRESSO global test for each BMI-associated metabolite. Overall, only 17 out of 324 (5.25%) metabolites reached nominal significance (*p* < 0.05) in the pleiotropy tests, and only two were in our causality groups (both unknowns: HILIC-pos_4106 and C8-pos_871). Thus, horizontal pleiotropy of BMI SNPs is unlikely to bias the majority of our metabolite classifications using G_B_. However, due to insufficient numbers of strong instruments available in our metabolite GWAS or in the literature, a comparable analysis could not be performed to assess pleiotropy in the metabolite instruments.
